# Uclacyanin proteins are required for lignified nanodomain formation within Casparian strips

**DOI:** 10.1101/2020.05.01.071738

**Authors:** Guilhem Reyt, Zhenfei Chao, Paulina Flis, Gabriel Castrillo, Dai-Yin Chao, David E. Salt

## Abstract

Casparian strips (CS) are cell wall modifications of vascular plants restricting extracellular free diffusion into and out the vascular system. This barrier plays a critical role in controlling the acquisition of nutrients and water necessary for normal plant development. CS are formed by the precise deposition of a band of lignin approximately 2 μm wide and 150 nm thick spanning the apoplastic space between adjacent endodermal cells. Here, we identified a copper-containing protein, Uclacyanin1 (UCC1) that is sub-compartmentalised within the CS. UCC1 forms a central CS nanodomain in comparison with other CS-located proteins that are found to be mainly accumulated at the periphery of the CS. We found that loss-of-function of two uclacyanins (*UCC1* and *UCC2*) reduces lignification specifically in this central CS nanodomain, revealing a nano-compartmentalised machinery for lignin polymerisation. This lack of lignification leads to increased endodermal permeability, and consequently to a loss of mineral nutrient homeostasis.

## Introduction

Plant roots perform the critical function of controlling the uptake of water and mineral nutrients from the soil essential for plant growth and development. A specialized cell layer in the root called the endodermis plays a key role in the selective uptake of mineral nutrients into the stele for translocation to the shoot [1–4]. Of vital importance to this function are Casparian strips (CS), which are belt-like lignin structures surrounding each endodermal cell, that interlock to form a barrier to diffusion in the apoplast [5,6]. This barrier is thought to enable the endodermis to exert control over uptake of water and solutes from the environment into the plant and perhaps also to control biotic interactions [7,8].

The precise deposition of lignin for CS formation requires a signalling pathway involving the kinases SGN1 and SGN3 controlling the spatial production of reactive oxygen species through the activation of NADPH oxidases [2,9–12]. Lignin polymerisation requires the localised action of a peroxidase (PER64) [9,13], and a dirigent-like protein (ESB1) [1]. This biosynthetic machinery is likely placed at the CS deposition site by association with CASPARIAN STRIP MEMBRANE DOMAIN PROTEINS (CASPs) [14]. The CASPs form a highly scaffolded transmembrane domain guiding where the Casparian strip forms. Furthermore, the receptor like kinase SGN3 acts as a sensor of CS integrity by inducing over-lignification of the endodermal cells when the CS is defective [2,15].

A network of transcriptional factors involving SHR, SCR and MYB36 controls endodermal differentiation [16,17]. The MYB36 transcription factor controls the expression of most of the described genes associated with CS formation, including *CASPs, ESB1* and *PER64* [3,18]. Characterisation of other MYB36-regulated genes could thus lead to the identification of new actors involved in CS formation.

Here, we identified a copper-containing protein, Uclacyanin1 (UCC1) among the MYB36-regulated genes. UCC1 reveals a nano-compartmentalisation of the machinery required for lignin polymerisation at the CS, where UCC1 occupies a central CS nanodomain in comparison with other CS-located proteins. The loss of function of two uclacyanins (*UCC1* and *UCC2*) leads to an atypical CS formation, where a lack of lignification is observed in this central CS nanodomain. This defect in lignification leads to increased endodermal permeability, and consequently to a loss of mineral nutrient homeostasis.

## Results and discussion

Among the genes downregulated in a *myb36* loss-of-function mutant [3,18], we identified a *Uclacyanin1* gene (*UCC1*), that belongs to the copper-containing phytocyanins family [19] (Figure 1A). This family is divided in three sub-families according to their copper binding amino acid: the uclacyanins, stellacyanins and plantacyanins. The functional role of these proteins remains unknown. However, biophysical and structural data of several phytocyanins suggest their implication in redox reactions with small molecular weight compounds [19–24]. The expression of several members of this family have been previously shown to be associated with lignified tissues [25–27]. We looked at the endodermal spatiotemporal expression pattern of the different members of this family (Figure 1A). We found that *UCC1, UCC2* and to a less extent *UCC8* and *STC1* are expressed in the endodermis similarly to that observed for *CASP1* and *ESB1*. Due to the lack of T-DNA insertional mutants in the *UCC1* gene, we generated two loss-of-function mutants, *ucc1.1* and *ucc1.2*, using CRISPR/Cas9 technology (Figure 1B). These mutants bear a single base deletion (*ucc1.1*) and an insertion (*ucc1.2*), leading to shifts in the reading frame in both. The two *ucc1* alleles show a slight although significant delay in formation of an apoplastic barrier, as visualized by the uptake of the apoplastic tracer propidium iodide (PI) after 10 min of staining (Figure 1C). The T-DNA mutant *ucc2.1* did not show an increase of PI permeability. Notably, in a double mutant *ucc1.2 ucc2.1*, a strong delay of PI permeability is observed similarly to that found in other CS mutants such as *esb1* and *casp1 casp3* (Supplementary Figure 1A). However, the number of cells permeable to PI in *ucc1.2 ucc2.1* is highly variable in comparison with *casp1 casp3* and *esb1*. When the incubation time with PI is increased to 20 min, *ucc1.2 ucc2.1* displays an enhanced permeability in comparison with *esb1*. Taken together, these results highlight the redundant role of UCC1 and UCC2 for establishing a functional apoplastic barrier. No additive effect was observed in the triple mutant *ucc1.1 casp1 casp3* or in the doubles *ucc1.2 esb1* and *ucc1.2 sgn3* in comparison with *casp1 casp3, esb1* and *sgn3* respectively (Supplementary Figure 1A). The loss of CS integrity in *esb1, myb36* and *casp1 casp3* is accompanied by an increased deposition of suberin [1–3] as a compensatory mechanism under the control of SGN3 [15]. We analysed the pattern of suberin deposition using fluorol yellow 088 staining in the *ucc* mutants (Figure 1D). Deposition of suberin lamellas did not increase in the single mutants *ucc1.1, ucc1.2* and *ucc2.1*. However, the double mutant *ucc1.2 ucc2.1* induces an increase in endodermal suberization compared with wild type plants (WT). This increase is not as strong as that observed in *esb1* or *casp1 casp3* (Supplementary Figure 1B). The combination of *ucc1* mutations in *esb1* and *casp1 casp3* do not affect the enhanced suberization of *esb1* or *casp1 casp3*.

**Figure 1.**
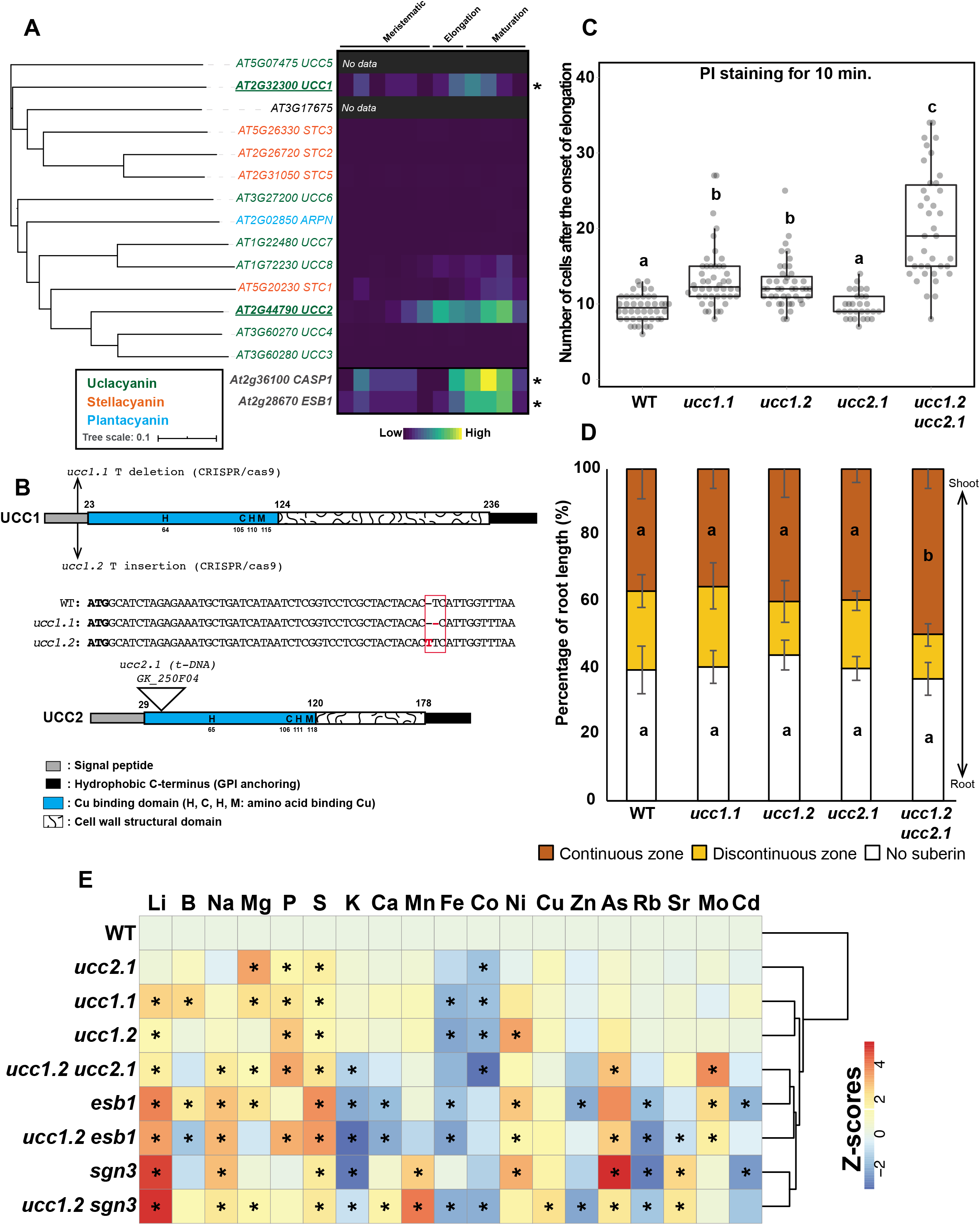
Uclacyanins UCC1 and UCC2 are required for a functional Casparian strip. **A.** (left) Figure shows a phylogenetic analysis of phytocyanins protein family in *A. thaliana*. The tree was built using the full-length amino acid sequences for all proteins. Different colours represent the three phytocyanins subfamilies: uclacyanins, stellacyanins and plantacyanins (39). In the tree, branch lengths are proportional to the number of substitutions per site. *AT3G17675* have been previously annotated as a stellacyanin (*STC4*), however the signal peptide for the secretion pathway and the hydrophobic extension for Glycosylphosphatidylinositol (GPI) anchoring are missing. (right) Heatmap showing the endodermal expression of the phytocyanins family in *A. thaliana* across the different root zones (Meristematic, Elongation, Maturation). For the analysis, expression data was collected from the Bio-Analytic Resource database, AtGenExpress Consortium. The expression of two endodermal localised proteins, CASP1 and ESB1, were added to the analysis as a reference. Asterisks indicate a significant downregulation in a *myb36* mutant according to [18]. B. Schematic representation of the UCC1 and UCC2 proteins showing the different protein domains and the types of mutations. Domains were defined according to [19]. **C.** Boxplot analysis showing the number of the cells from the onset of elongation permeable to propidium iodide in wild type plants (WT), *ucc1* mutants (*ucc1.1* and *ucc1.2), ucc2* mutant (*ucc2.1*), and the double mutant *ucc1.2 ucc2.1*. Data were collected from two independent experiments (n ≥ 29). Different letters represent significant differences between genotypes in a Mann-Whitney test (*p* < 0.01). **D.** Diagram shows the quantification analysis of the endodermal suberization in the plant genotypes used in C. Each colour in the graph represents the percentage of the root length (percentage of root length (%)) that is unsuberised (white), discontinuously suberised (yellow), continuously suberised (orange). Suberin was staining with Fluorol yellow 088. (n ≥ 18). Error bars in the figure are the standard deviation (SD). Different letters represent significant differences between genotypes using a Mann-Whitney test (*p* < 0.01). **E.** Heatmap representing the io-nomic profiles (z-scores) of wild type plants (WT), and a collection of mutants with a defective casparian strip: *ucc2.1, ucc1.1, ucc1.2, ucc1.2 ucc2.1, esb1, ucc1.2 esb1, sgn3* and *ucc1.2 sgn3* grown in full nutrient conditions on agar plate for 2 weeks (n=10). Elements concentration were determined by ICP-MS and the raw data is available in the Supplementary Table 1. Asterisks indicate a significant difference in comparison with WT using a *t*-test (*p*<0.01). Columns (Genotypes) were subjected to hierarchical clustering analysis.

Disruption of CS is known to affect the composition of the leaf ionome as observed in *esb1, myb36, casp1 casp3, sgn3* and *lotr1* [1–3,28]. In order to determine the contribution of UCC1 and UCC2 in maintaining mineral nutrient homeostasis, we analysed the leaf ionome in different mutant combinations (Figure1E, Supplementary figure 1C, Supplementary table 1) using inductively coupled plasma mass spectrometry (ICP-MS). The *ucc2.1* mutant only exhibits minor ionomic changes and is similar to WT in a principal component analysis (PCA; Supplementary figure 1C). The single *ucc1* and the double *ucc1.2 ucc2.1* mutants present a larger number of elemental changes in the leaf ionome in comparison with *ucc2.1*. Moreover, the leaf ionome of *ucc1.2 ucc2.1* shows similarities to the *esb1* ionome in a hierarchical clustering analysis and in a PCA. Adding the *ucc1.2* mutation into a *sgn3* mutant background does not strongly affect the leaf ionome in comparison with *sgn3*. The same effect but to a lesser extent is observed for *ucc1.2 esb1* in comparison with *esb1* (Supplementary figure 1C). The higher endodermal permeability and the atypical ionomic profile observed (Figure 1 and Supplementary figure 1) strongly suggest that UCC play an important role in CS formation. To identify the cell type in which the *UCC1* promoter is activated, we fused the *UCC1* promoter to *GFP* (Supplementary figure 2A). GFP was accumulated only in the endodermis, from the elongation zone and further up into the zone of differentiated endodermal cells. According to the literature, the phytocyanin proteins are predicted to be located on the cell surface and anchored to the plasma membrane via a glycosylphosphatidylinositol (GPI) anchor [29]. Several members of this family, including UCC2, have been shown to be GPI-anchored using a proteomic approach [29]. In order to determine the precise UCC1 localization, we generated a line (*pUCC1::mCherry-UCC1*) expressing a tagged UCC1 (Figure 2A). The mCherry-UCC1 fusion accumulates at the endodermal cell junctions where the CS is located (Figure 2B). It first appears in a discontinuous manner in the early stage of endodermal differentiation, and then forms a more continuous band later in the endodermis development. Highly mobile vesicles associated with mCherry-UCC1 are also seen in the endodermis (Figure 2B, Supplementary video 1).

**Figure 2.**
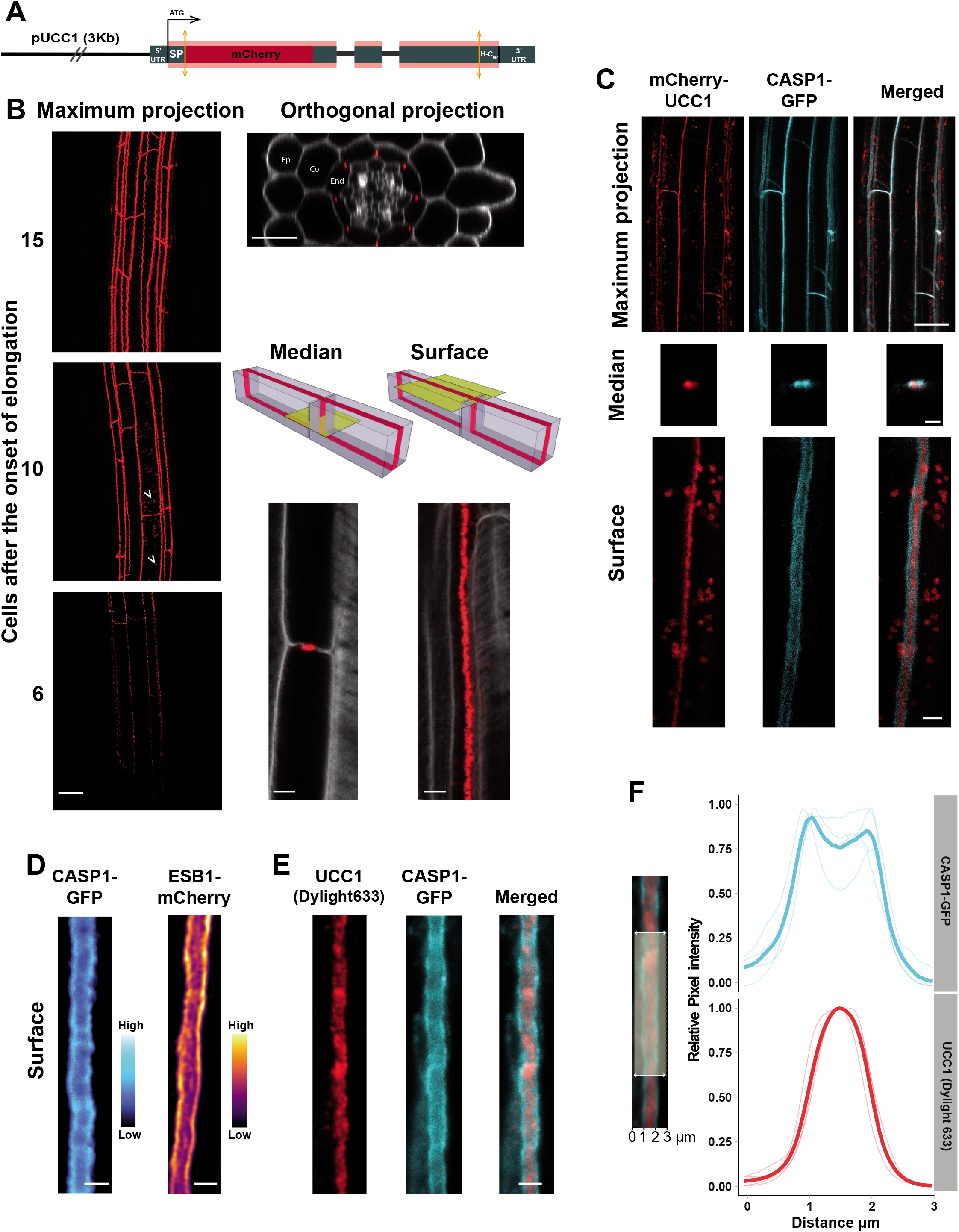
UCC1 defines a new central sub-domain in the Casparian strip. **A.** Diagram representing the construct *pUCC1::mCherry-UCC1* (UTR: Untranslated region, SP: Signal peptide, H Cter: Hydrophobic C-terminus for GPI anchoring). **B.** Maximum intensity projection, orthogonal, median and surface views of confocal sections of plants expressing *pUCC1::mCherry-UCC1* (red) in cleared roots. In the case of maximum intensity projection (Maximum projection) figure represents different regions of the root measured as number of cells after the onset of elongation. White arrow heads point to vesicles containing *mCherry-UCC1*. For the orthogonal, median and surface views, cell walls were stained with Calcofluor white (grey in the figures). Scale bar = 20 μm for the maximum projection and orthogonal views. Scale bar = 5 μm for the median and surface views (Ep: epidermis, Co: cortex, End: endodermis). **C.** Maximum intensity projection, median and surface view of confocal sections of plants expressing *CASP1-GFP* (cyan) and *mCherry-UCC1* (red). Signal was capture at the 10^th^endodermal cell after the onset of elongation observed *in vivo*. Scale bar = 20 μm for maximum projection and 3 μm for median and surface view. **D.** *In vivo* observation of the surface view of an endodermal cell expressing *pESB1::ESB1-mCherry* or *pCASP1::CASP1-GFP*. Scale bar = 2 μm. **E.** Immunolocalization assay of UCC1 protein (red) in plant expressing*pCASP1::CASP1-GFP* (cyan). A primary polyclonal antibody targeting UCC1 was used in combination with a secondary antibody conjugated with Dylight 633. Scale bar = 2 μm. **F.** Graph presenting the distribution of normalised pixels intensity (Relative Pixel Intensity, 0 to 1) across the Casparian strip (Distance μm) for *CASP1-GFP* fluorescence (cyan) and *UCC1* immunofluorescence (red, Dylight 633). Light curves represent individual replicates coming from individual plants (n = 4). Each replicate is the average pixel intensity across a segment of 25 μm along the Casparian strip axis. Dark curves represent the mean values for *CASP1-GFP* and *UCC1* immunofluorescence.

When we analysed the localization of mCherry-UCC1 in plants expressing *CASP1-GFP*, we observed a similar pattern of localization at the CS as seen in the maximum projection view of endodermal cells in Figure 2C. However, at higher magnification, in both the median and surface views we observed mCherry-UCC1 to occupy a more central position in comparison with CASP1-GFP. This was further confirmed using super-resolution structure illumination microscopy (Supplementary figure 2B). Moreover, CASP1-GFP does not form a homogeneous domain, but is found to accumulate more at the periphery of the CS and less in the centre where UCC1 is observed. Subsequently, we checked if mCherry-UCC1 colocalises with lignin deposition in the CS (Supplementary figure 2C). At the early stage of endodermal differentiation, lignin presents a nearly perfect colocalization with mCherry-UCC1. However, the line expressing mCherry-UCC1 also presents ectopic lignification later in development. This ectopic lignification is similar to that observed in other CS mutants such as *casp1 casp3* and *esb1* [1,14]. Expression of *mCherry-UCC1* also delayed the formation of a functional barrier to PI and promotes a significant increase of suberin deposition (Supplementary figure 2D-F). These observations taken together show that expressing *UCC1* with a fluorescent tag causes disruption of the CS.

To establish if the expression of *mCherry-UCC1* can affect the pattern of accumulation of CASP1-GFP presented in Figure 2C and Supplementary figure 2B, we examined plants expressing *CASP1-GFP* only (Figure 2D). CASP1-GFP is still more accumulated at the periphery, and this was also observed for another member of the CS machinery ESB1-mCherry. Immunolocalization confirmed that native UCC1 protein is specifically localized to the CS (Figure 2E), and its localization was strongly reduced in *ucc1.2* and *ucc1.2 ucc2.1* mutants, and abolished in a *myb36* mutant (Supplementary figure 2G). Immunolocalization confirmed our previous observations of a central localization of UCC1 in the CS as compared to CASP1-GFP (Figure 2E). Quantification of pixel intensity across the CS reveals that UCC1 is highly accumulated in the middle of the CS where CASP1-GFP is slightly less accumulated compared to the periphery of the CS domain (Figure 2F). The CASP1 domain of the plasma membrane is defined as a microdomain [30] as its width is around 1.5μm. UCC1 with a more central accumulation has a width equal or slightly smaller than 1μm, UCC1 can then be classified as a nanodomain of the CS. This subdomain structure has not been previously reported. However, previous studies for CASP1, ESB1 and Peroxidase64 (PER64) [14,31] do appear to show an enrichment of these proteins at the periphery of the CS. This means that the level of organisation observed at the plasma membrane for CASP1 is conserved for cell wall proteins such as ESB1 and PER64. To date, UCC1 is the only protein found in the central nanodomain of the CS, revealing a new level of internal structure within the CS.

To characterise the formation of the UCC1 and CASP1 subdomains, we tracked endodermal differentiation cell-by-cell. CASP1-GFP and mCherry-UCC1 are found to be accumulated in the endodermis concomitantly between 4 and 6 cells after the onset of elongation, at the periphery of the cells and not yet at the CS (Figure 3 A-B) which is consistent with a common transcriptional regulation by MYB36. We then tracked the central accumulation of CASP1-GFP and mCherry-UCC1 at the CS. We observed that the localization of CASP1-GFP and mCherry-UCC1 at the CS mainly occurs at the eighth endodermal cell after the onset of elongation (Figure 3B). Although, mCherry-UCC1, but not CASP1-GFP, was observed to be centrally localised in several independent events at the seventh cell after the onset of elongation. This suggests that CASP1 localization at the CS is not required for the recruitment of UCC1. This was further confirmed by the targeting of UCC1 at the CS in a *casp1 casp3* mutant (Figure 3C and Supplementary figure 3). However, UCC1 is not able to form a continuous domain in *casp1 casp3* as in WT. This was also observed in the *esb1, sgn3* and *esb1 sgn3* mutants. We then tested the reciprocity to know if UCC1 and UCC2 are required for CASP1-GFP localization (Figure 3D). In the *ucc1.1* and *ucc1.1 ucc2.1* mutants, CASP1-GFP is able to localise at the CS domain and form a continuous domain without disruption. This demonstrate that UCC1 and UCC2 are not required for CASP1 localization to the CS domain. However, the absence of UCC1 and UCC2 does affect CASP1-GFP localization at a nanoscale resolution (Figure 3E). The exclusion of CASP1-GFP from the central nanodomain of the CS is reduced in the *ucc1* mutants and tends to disappear in the double *ucc1.1 ucc2.1* mutant. This indicates a role for UCC1 and UCC2 in the formation of the central CS nanodomain.

**Figure 3.**
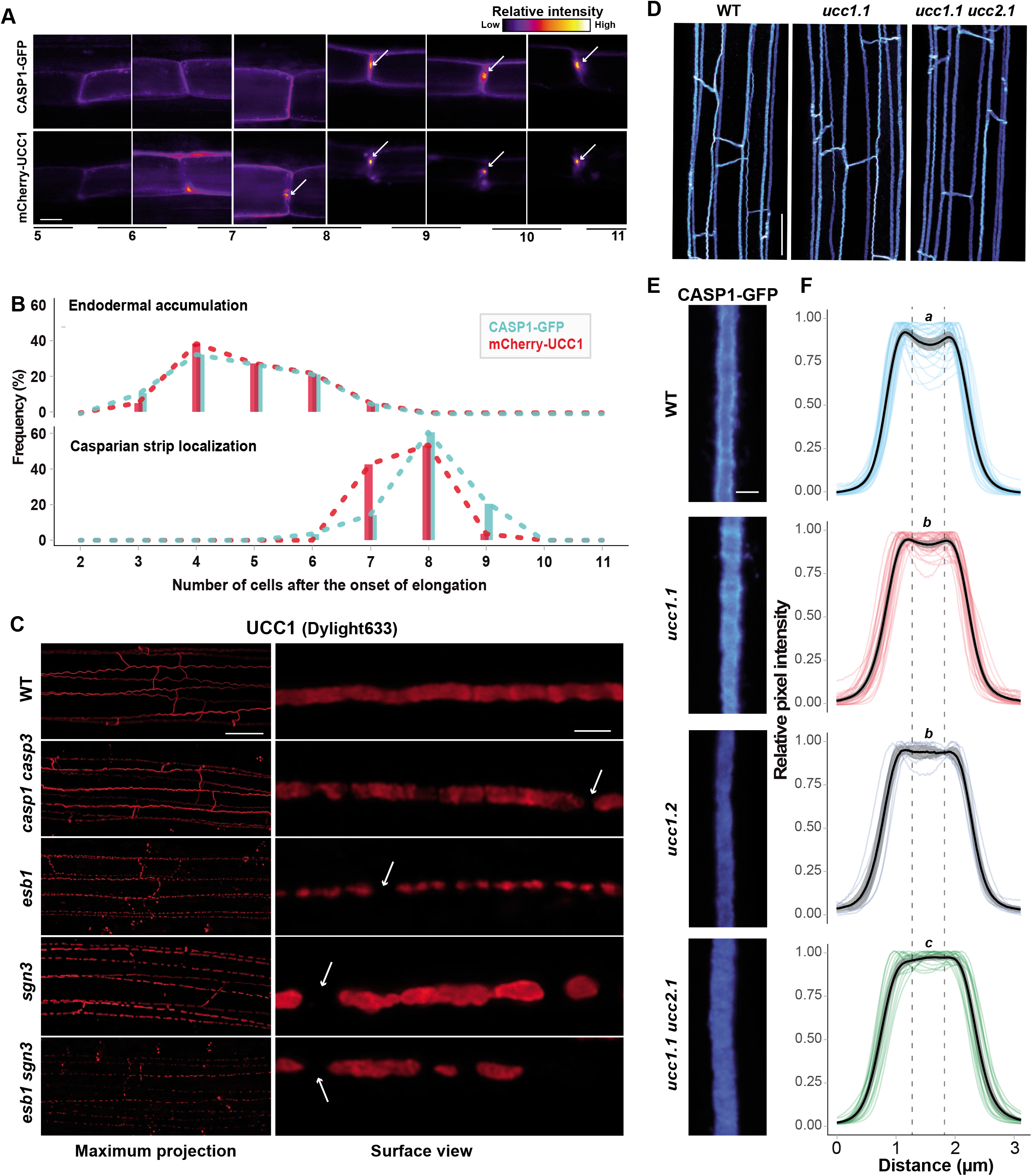
Relations between UCC1 positioning and other components of the Casparian strips machinery. **A.** Analysis of the spatial distribution of CASP1 and UCC1 at the endodermal cell junctions. Images were generated from the same plant co-expressing CASP1-GFP and mCherry-UCC1 using confocal microscopy. The numbers at the bottom of the figure indicate the number of cells after the onset of elongation. White arrows indicate the central accumulation for CASP1-GFP or mCherry-UCC1. Scale bar = 6 μm. **B.** Histograms showing the frequency distribution (Frequency (%)) of the onset of expression (upper plot, n = 18) and the onset of localization at the Casparian strip of CASP1-GFP and mCherry-UCC1 (lower plot, n = 28). **C.** (left panel) Maximum intensity projection and (right panel) surface view of UCC1 immunolocalization (red) at 10 cells after the onset of elongation in wild type plants (WT) and a collection of Casparian strips mutants: *casp1 casp3, esb1, sgn3, esb1 sgn3*. White arrows show gaps in the UCC1 localization. Scale bar = 20 μm for the maximum projections and 2 μm the surface views. **D.** Maximum intensity projection of CASP1-GFP localization in cleared root of wild type plants (WT), and the mutants *ucc1.1* and *ucc1.1 ucc2.1*. Scale bar = 20 μm. **E.** Surface view of the localization of CASP1-GFP in cleared root of wild type plants (WT), and the mutants: *ucc1.1, ucc1.2* and *ucc1.1 ucc2.1*. Scale bar = 2 μm. **F.** Quantification of normalised pixels intensity (Relative Pixel Intensity; 0 to 1) across the Casparian strip in plants expressing CASP1-GFP. The plots are showing the intensity profile for individual replicates (n ≥ 10), the mean value (black line) and the 95% confidence interval (grey interval). Each replicate corresponds to the quantification of one picture containing a Casparian strip segment of approximately 25μm long. The pictures were generated at the 15^th^ cells after the onset of elongation from at least 8 individual plants per genotype. Intensity profiles across the Casparian strip were always measured in the same orientation, from the cortical side toward the pericycle side of the endodermis. Letters indicate statistically significant differences between genotypes for the intensity values comprised between the dashed lines using an ANOVA and Tukey’s test as *post hoc* analysis (*p*<0.01).

Knowing that UCC1 localizes at the CS (Figure 2C-F) and mutations in *UCC1* and *UCC2* cause a strong defect in root apoplastic permeability (Figure 1C, Supplementary figure 1A), we looked at lignin deposition at the CS in *ucc* mutants (Supplementary figure 4A). Mutations in *UCC1* and/or *UCC2* do not cause an obvious disruption in the CS as observed in *casp1 casp3, esb1, sgn3* (Supplementary Figure 4A), and other previously identified CS mutants, where clear gaps can be observed [1,2,7,10,12–14,28,32]. Surprisingly thought, mutations in *UCC1* do reduce the amount of lignin in the central nanodomain of the CS, as observed using confocal imaging (Figure 4A), and super-resolution structure illumination microscopy (Supplementary figure 4B). Pixel quantification across the CS reveals that *ucc1* mutants show a lower lignification in the central nanodomain (Figure 4B) where UCC1 accumulates (Figure 2C-F). This decrease is not observed in *ucc2.1*. However, a further decrease is observed in the double mutant *ucc1.2 ucc2.1* in comparison with *ucc1* mutants. This is consistent with the increased root permeability observed in *ucc1.2 ucc2.1* in comparison with the single mutants (Figure 1C, Supplementary figure 1A). This reveals that CS permeability can be strongly affected in the absence of clear gaps in the CS. Furthermore, we observed ectopic lignification on the cortical side of the endodermis on a few occasions in the *ucc1* mutants, and at a higher frequency in the double mutant *ucc1.2 ucc2.1* (Figure 4A-B, Supplementary figure 4). This is a typical phenotype, along with increased suberin deposition, that is due to the SGN3-dependent compensatory mechanism observed in most mutants with defective CS [1,2,14,18,33]. However, the ectopic lignification and enhanced suberization are observed to a lesser degree in *ucc1.2 ucc2.1* in comparison with other CS mutants such as *casp1 casp3* and *esb1* mutants (Figure 1D and 4, Supplementary figure 1B and 4). This could be explained by a more conditional leakiness of the CS in *ucc1 ucc2* mutants. Discontinuities below the resolution of light microscopy could occur in *ucc1 ucc2* mutants. This could lead to a full permeability for low molecular weight compounds such as ions and PI but an intermediate permeability for high molecular weight compounds such as the CIF peptides required for ectopic lignification and enhanced suberization [10,32].

**Figure 4.**
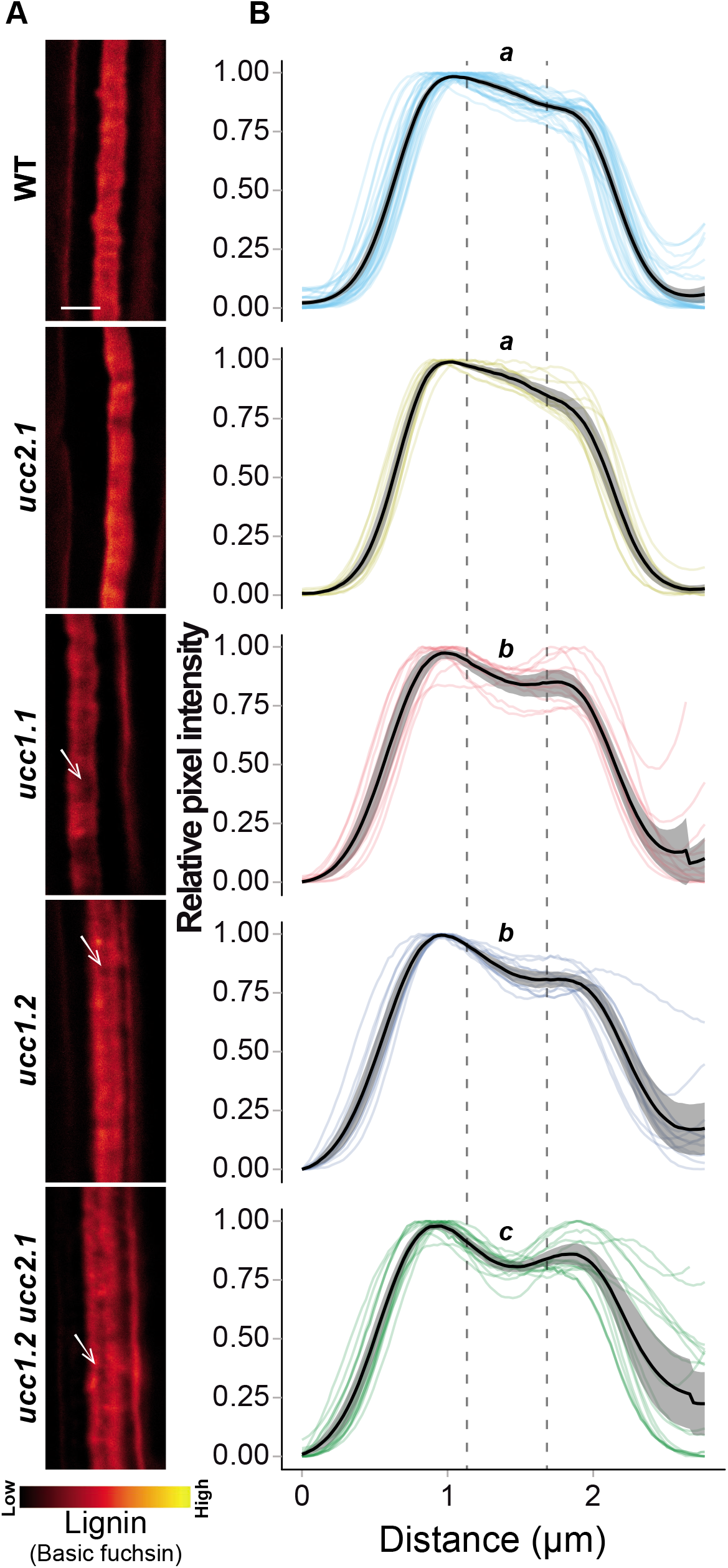
UCC1 and UCC2 are necessary for the central lignification of the Casparian strip. **A.** Surface view of the Casparian strip lignin stained with Basic fuchsin in wild type plants (WT), and the mutants *ucc2.1, ucc1.1, ucc1.2* and *ucc1.2 ucc2.1*. Whites arrows show lack of lignification in the central domain of the Casparian strip across the different genotypes. Scale bar = 2 μm. **B.** Quantification of normalised pixels intensity (0 to 1) (Relative pixel intensity) across the Casparian strip using surface views as shown in A. The plots show the intensity profile for individual plants (n ≥ 13). In the figure, the mean value is represented by a black line and the 95% confidence interval is in grey. The data were generated using individual pictures containing a Casparian strip segment of approximately 25 μm long. The pictures were taken at the 15^th^ cells after the onset of elongation and the intensity profiles were measured in the same direction, from the cortex side toward the pericycle side of the endodermis. Letters indicate statistically significant differences between genotypes using an ANOVA and Tukey’s test as *post hoc* analysis (*p*<0.01).

In conclusion, this study reveals the first loss-of-function phenotype for members of the plant-specific blue copper protein family of phytocyanins. Several studies suggested their implication in lignin polymerisation [25–27], but this has never previously been shown. Our analysis indicates a role for the Uclacyanins in the deposition of lignin in a newly discovered nanoscale domain within the CS. Further, the subcellular localization of UCC1, and the phenotype of the *ucc*(*s*) mutants, reveals a sub-compartmentalisation of the machinery required for lignin polymerisation at the CS.

## Author contributions

Conceptualization, G.R., G.C. and D.E.S.; Formal Analysis, G.R.; Investigation, G.R., Z.C., P.F. and G.C.; Methodology G.R. and Z.C.; Writing – Original Draft Preparation, G.R.; Writing – Review & Editing G.R., Z.C., P.F., G.C., D-Y.C. and D.E.S.; Supervision, D.E.S.

---

**Supplementary figure 1.**
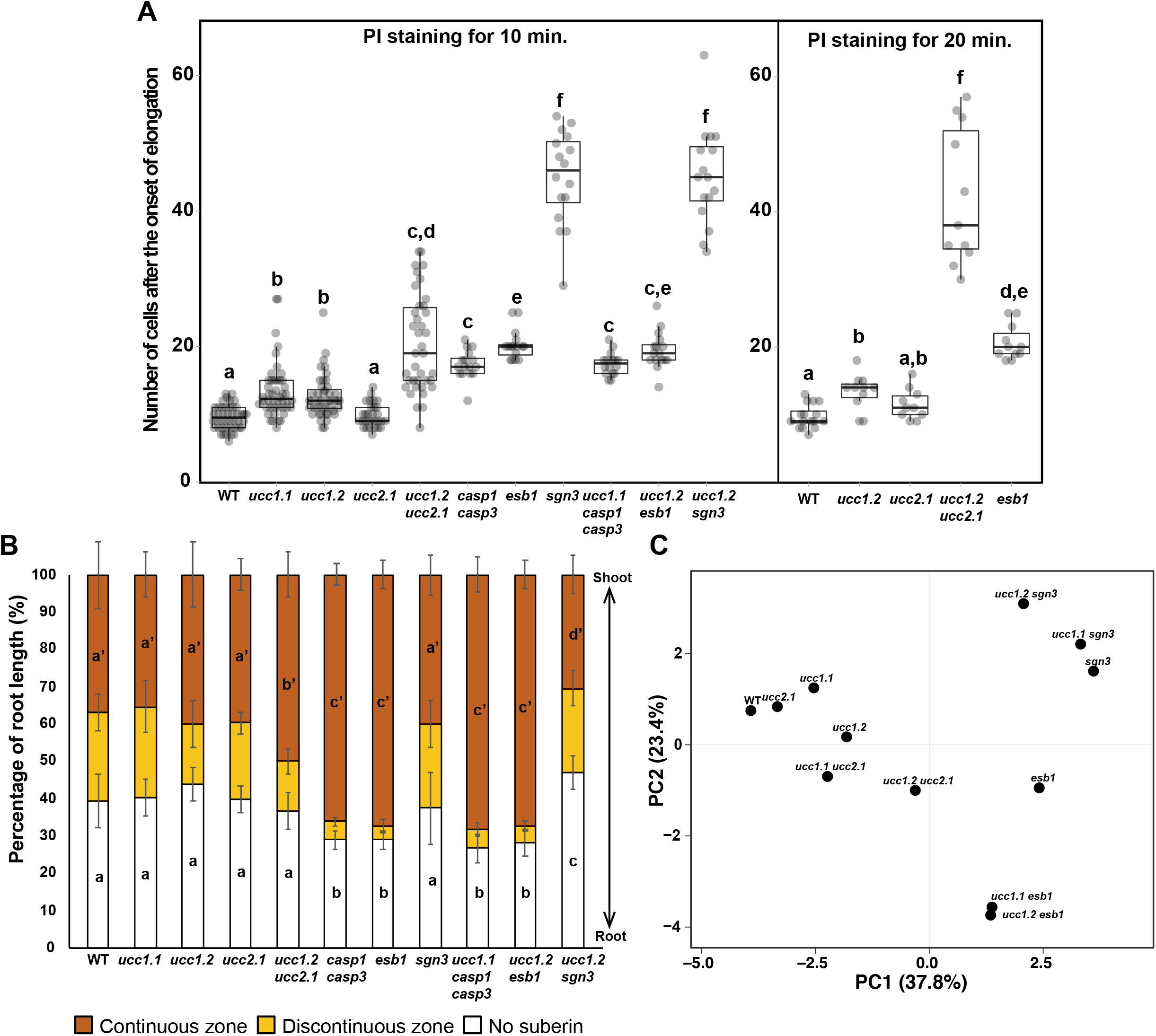
Uclacyanins UCC1 and UCC2 are required for the correct function of the Casparian strips. **A.** Boxplot showing the number of cells, from the onset of elongation, permeable to propidium iodide (PI) after 10 min or 20 min of root staining in wild type plants (WT) and a collection of mutants (*ucc1.1, ucc1.2, ucc2.1, ucc1.2 ucc2.1, casp1 casp3, esb1, sgn3, ucc1.1 casp1 casp3, ucc1.2 esb1, ucc1.2 sgn3*) The number of plants analysed was n ≥ 16 and n ≥ 10 for the 10 min and 20 min of PI exposition, respectively. Different letters represent significant differences between genotypes using a Mann-Whitney test (*p* < 0.01). **B.** Quantification of the endodermal suberization using the suberin specific staining Fluorol yellow 088 in the genotypes from A. The results (Percentage of root length (%)) represent the percentage of the root that remains unsuberised (white), discontinuously suberised (yellow) or continuously suberised (orange). In all cases the number of plants analysed was n ≥ 15, error bars mean standards deviation (SD). Different letters represent significant differences between genotypes using a Mann-Whitney test (*p* < 0.01). **C.** Principal component analysis (PCA) of the shoot ionomic profiles of wild type plants (WT) and the defective CS mutants *ucc2.1, ucc1.1, ucc1.2, ucc1.1 ucc2.1, ucc1.2 ucc2.1, esb1, ucc1.1 esb1, ucc1.2 esb1, sgn3, ucc1.2 sgn3* and *ucc1.1 sgn3* grown in agar plates. The PCA was performed using the average of the elemental profiles from each genotype (n=10) available in the Supplementary Table 1.

**Supplementary figure 2.**
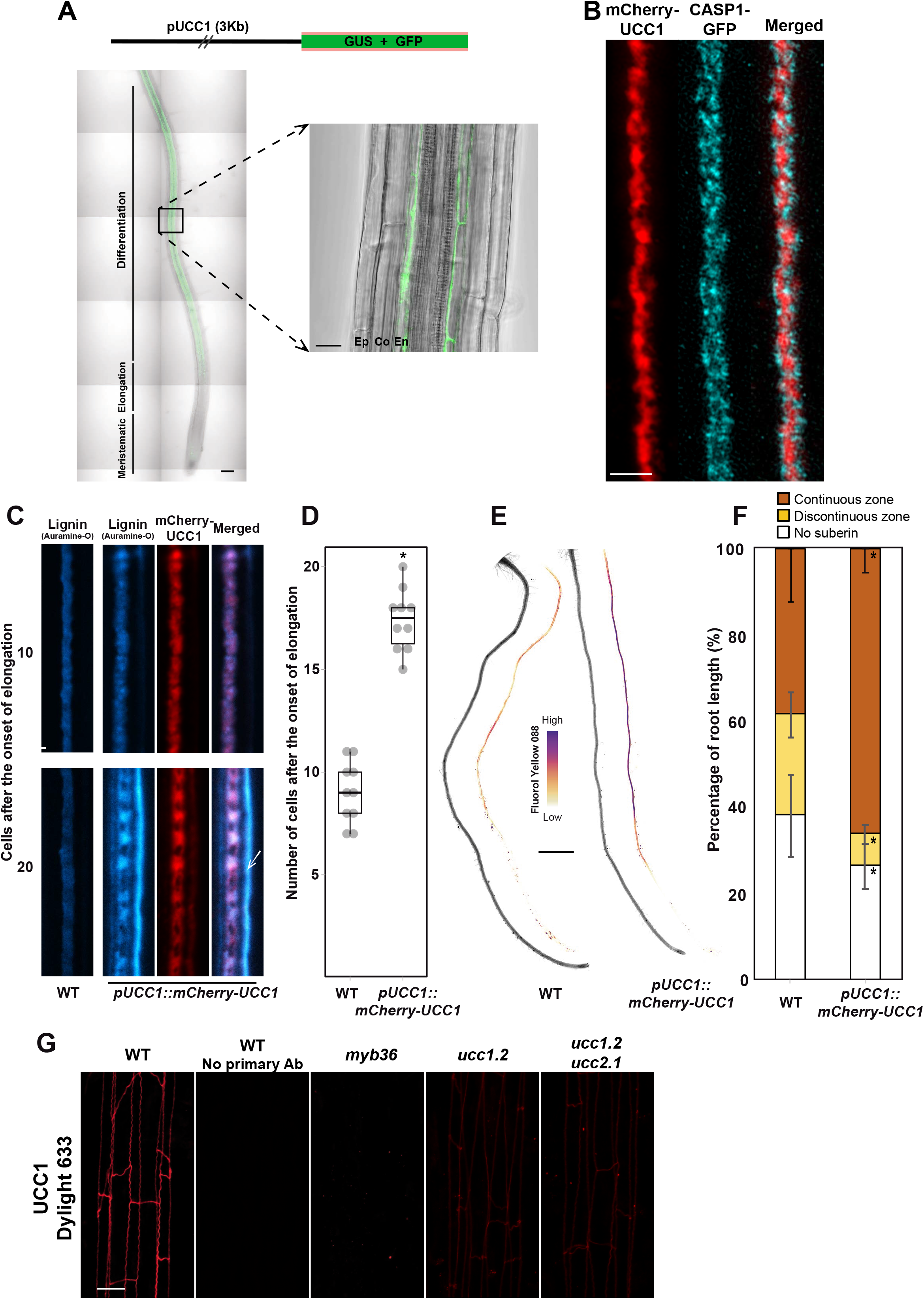
UCC1 accumulates in the root endodermis and defines a central sub-domain at the Casparian strip. **A.** (Top) Schematic representation of the construct used to study the expression of *UCC1* in the root. Approximately 3 Kb of the promoter region of the gene *UCC1* (pUCC1 (3Kb)) was used to drive the expression of the β-glucuronidase (GUS) and the green fluorescent protein (GFP). (Bottom) Pictures showing the GFP accumulation in the root of plants expressing the construct *pUCC1::GUS-GFP*. The fluorescence (GFP, green) and brightfield (grey) pictures are merged. The different zones of the root (meristematic, elongation and differentiation) are highlighted in the figure. Scale bars = 100 μm (left) and 20 μm (right). (Ep: Epidermis; Co: Cortex; En: Endodermis). **B.** Structure illumination microscopy of a surface view of plants expressing CASP1-GFP (cyan) and mCherry-UCC1 (red) at the 10^th^ endodermal cell after the onset of elongation in cleared root. Scale bar = 2 μm. **C.** Surface view of Casparian strip lignin stained with Auramine-O (cyan) in wild type plants (WT) and in a line expressing the construct *pUCC1::mCherry-UCC1* (red) in cleared roots at 10^th^ and 20^th^ cells after the onset of elongation. In the figure the white arrow shows ectopic lignification. Scale bar = 2μm. **D.** Boxplot showing the number of the cells from the onset of elongation permeable to propidium iodide in wild type plants (WT) and a line expressing *pUCC1::mCherry-UCC1*. For both genotypes 10 plants were analysed. Asterisk indicates a significant difference with WT using a Mann-Whitney test (*p* < 0.01). **E.** Brightfield and suberin staining using Fluorol yellow 088 of roots in WT and a line expressing *pUCC1::mCherry-UCC1*. Scale bar = 2 mm. **F.** Quantification of suberin staining with Fluorol yellow 088 along the root. The results are expressed in percentage of root length that is unsuberised (white), discontinuously suberised (yellow), and continuously suberised (orange). In all cases n ≥ 10 plants were used, error bars: SD. Asterisks indicates significant differences in comparison with WT for the same zone using a Mann-Whitney test (*p* < 0.01). G. The anti UCC1 antibody generated in this work is functional and specifically recognises the UCC1 protein. Maximum intensity projection of UCC1 immunolocalization (red) in wild type plants (WT), the mutants *myb36, ucc1.2, ucc1.2 ucc2.1* in the root at the 10^th^ cell after the onset of elongation. As a control the primary polyclonal antibody anti UCC1 was used or not (“No primary Ab”) in combination with a secondary antibody conjugated with Dylight 633. Scale bar = 20 μm.

**Supplementary figure 3.**
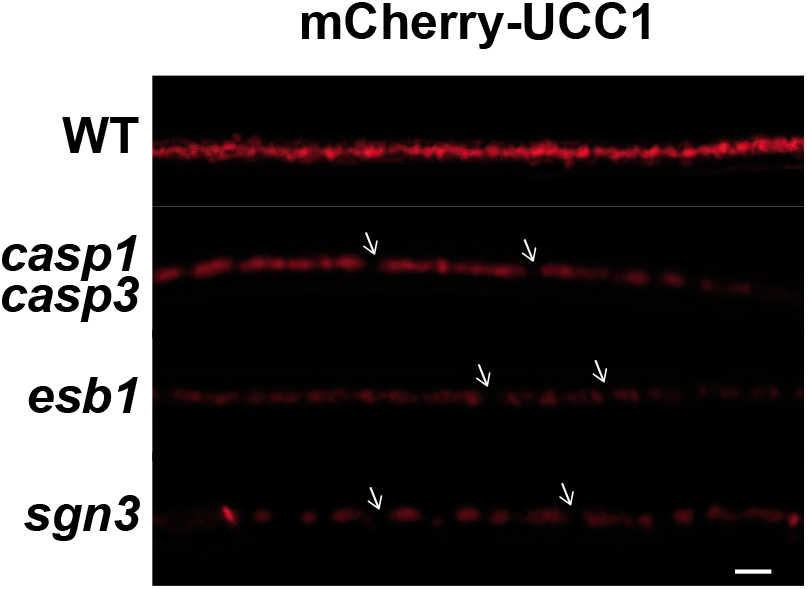
Other Casparian strip molecular players are required for UCC1 to fuse into a continuous band. Surface view of mCherry-UCC1 at the 15th cell after the onset of elongation in cleared root of WT, *casp1 casp3, esb1* and *sgn3*. White arrows represent discontinuous mCherry-UCC1 localization. Scale bar = 2 μm.

**Supplementary figure 4.**
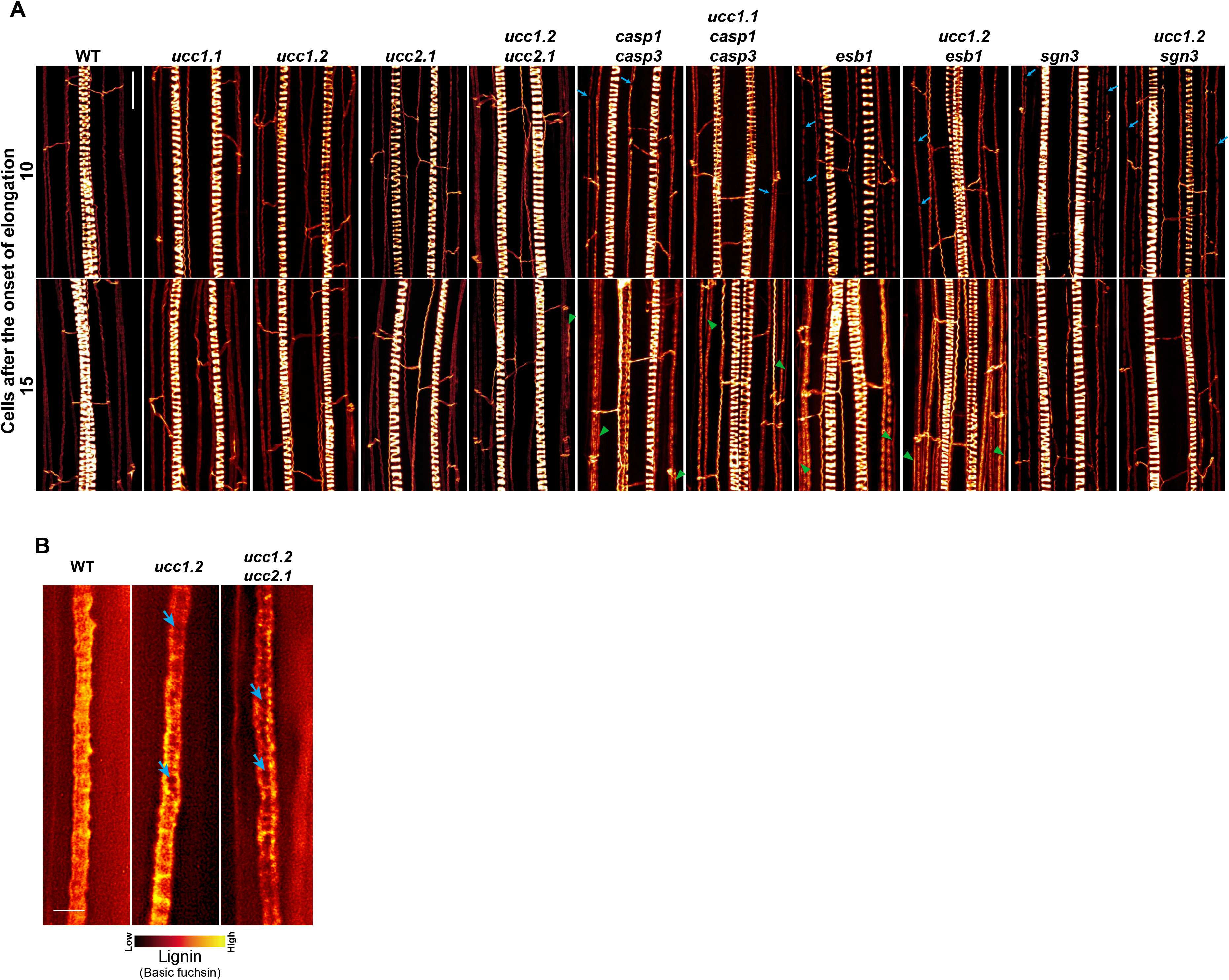
UCC1 and UCC2 are necessary for the central lignification of the Casparian strip. **A.** Figures show the maximum intensity projection of confocal sections of plant roots stained with basic fuchsin for visualising lignin. Pictures were taken using cleared roots at the 10th and 15th cells after the onset of elongation of wild type plants (WT), and a collections of mutants: *ucc1.1, ucc1.2, ucc2.1, ucc1.2 ucc2.1, casp1 casp3, ucc1.1 casp1 casp3, esb1, ucc1.2 esb1, sgn3* and *ucc1.2 sgn3*. Spiral structures in the centre of the root are the xylem. Blue arrows indicate gaps in the lignin deposition at the Casparian strip. Green triangles show ectopic lignification. Scale bar = 20 μm. **B.** Structure illumination microscopy of a surface view of Casparian strip lignin stained with basic fuchsine in cleared roots of WT, *ucc1.2* and *ucc1.2 ucc2.1*. Blue arrows show lack of lignification in the central domain of the Casparian strip. Scale bar = 2 μm.

**Supplementary table 1.**
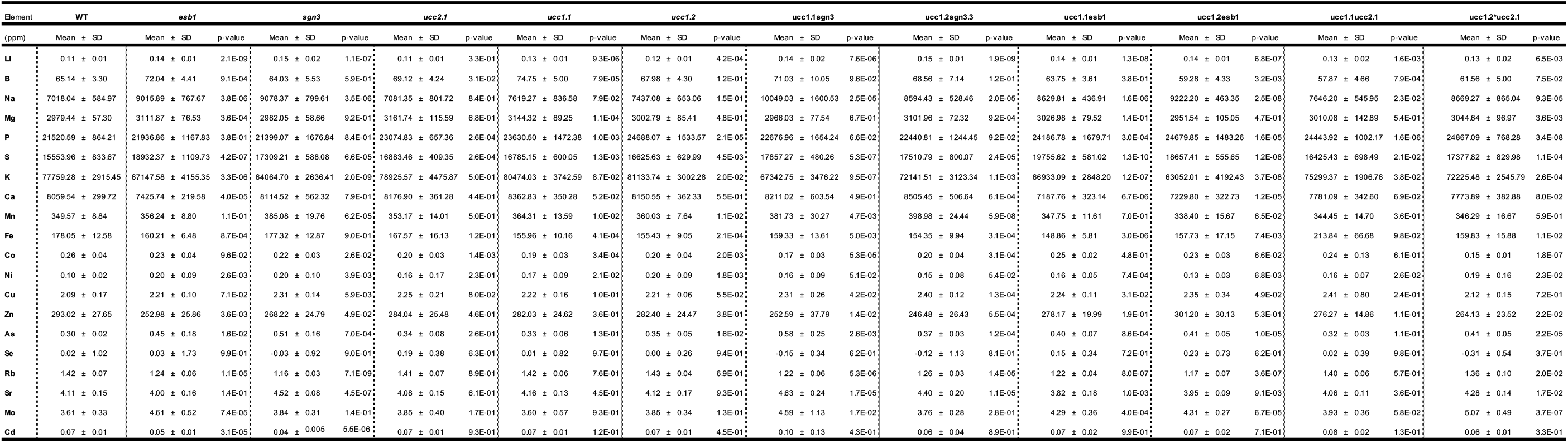
UCC1 and UCC2 are required for maintaining ion homeostasis in the shoot. Elemental content in shoot of *ucc2.1, ucc1.1, ucc1.2, ucc1. 1 ucc2.1, ucc1.2 ucc2.1, esb1, ucc1.1 esb1, ucc1.2 esb1, sgn3, ucc1.2 sgn3* and *ucc1.1 sgn3* mutants compared to WT grown in agar plates (long day, n=10). Elements concentration were determined by ICP-MS. Data are presented as mean ± standard deviation (SD). *t-tests* were performed to determine the significant differences to WT and the corresponding *p*-values are presented.

**Supplementary Video1.**

Time-lapse video showing vesicles of mCherry-UCC1 in the endodermis of *pUCC1::mCherry-UCC1*.

## Materials and methods

### Plant material

*Arabidopsis thaliana* ecotype Columbia (Col_0) and the following mutants and transgenic lines were used in this study: *sgn3 (sgn3.3;* SALK_043282[2]), *myb36 (myb36.2;* GK-543B11 [3]), *casp1 casp3 (casp1.1 casp3.1*, [14]), *pCASP1::CASP1-GFP* [14], *pESB1::ESB1-mCherry* [1], *ucc2.1* (GK_250F04).

The corresponding gene AGI are: *UCC1*, AT2G32300; *UCC2*, AT2G44790;

*SGN3*, At4g20140; *MYB36*, At5g57620; *CASP1*, At2g36100; *CASP3*, AT2G27370 *ESB1*, At2g28670.

### Generation of transgenic lines and CRISPR/Cas9 mutants

The line *pUCC1::mCherry-UCC1* was obtained using Gateway Cloning Technology (Invitrogen). A genomic DNA fragment containing the *UCC1* promoter (−3083bp before ATG), the UCC1 5’UTR and the UCC1 signal peptide (+78bp after ATG) was amplified by PCR using the following primers: F: GGGGACAACTTTGTATAGAAAAGTTG**GTTGAATTTCG-TAAGAGTTAGG** and R: GGGGACTGCTTTTTTGTACAAACTTGC**ATGGTCAG-TAGCTACTGTTAAACC**); and cloned in a pDONR P4-P1R. A second fragment containing the rest of the genomic UCC1 sequence (from +79 to +1081 from ATG) was amplified by PCR with the primers: F: GGGGACAGCTTTCTTGTACAAAGTGGTA**ACCATT-GGTGGTCCTAGTGGTTGG** and R: GGGGACAACTTTGTATAATAAAGTT-GA**CCCATATAAATTGTAATAATGTATTATAAAC** and cloned in a pDONR P2R-P3. Both fragments and a pEN-L1-mCherry-L2 vector were assembled in the expression vector pB7m34GW,3 [34]. The construct *pUCC1::GFP-GUS* was generated using Gateway cloning technology. The *UCC1* promoter (−3083bp to −83bp before ATG) was amplified by PCR with these primers: F: GGGGACAAGTTTGTACAAAAAAGCAGGCT**GTTGAATTTCGTAA-GAGTTAGG; Primer R :** GGGGACCACTTTGTACAAGAAAGCTGGGT**ATGACAT-ATGGTGTCAAATGTGTG** and cloned in pBGWFS7 [35]. Following Agrobacterium strain GV3101 transformation with the resulting vector, transgenic plants were generated by floral dipping [36].

The *ucc1.1* and *ucc1.2* mutants were obtained using CRISPR/Cas9 according to [37]. Two sgRNA targeting *UCC1* coding sequence (GTCCTCGCTACTACACTCA and GGTCCTAG-TGGTTGGACTG) were inserted in to the pHEE401 vector following Agrobacterium strain GV3101 transformation with the resulting vector, transgenic plants were generated by floral dipping [36]. Two individual homozygous lines (*ucc1.1*; *ucc1.2*) were identified at the generation T1.

### Growth Conditions

All seeds were surface sterilised, and then stratified for two days at 4°C. Seeds were directly germinated on plates containing ½ MS medium (Murashige and Skoog, Sigma) solidified with 0.8% agar, pH 5.7, and grown in a vertical position in a growth chamber under long-day conditions (16h light 22°C/8h dark 19°C, light intensity 100μE). Seedlings were analysed at 6-day-old for microscopy analysis, and 2-week-old for ionomic analysis.

### Phylogeny

A BLASTP was performed using UCC1 amino acid sequence against the Araport11 protein sequences dataset [38]. Only the matches with an Expect value (E-value) lower than 0.05 were considered. A first alignment was performed to select the protein sequence containing the amino acid required for copper binding as described in [19]. Genes were named and classified as stellacyanin, uclacyanin and plantacyanin according to the amino acid binding copper as described in [39]. A second alignment and tree assembling were performed using Clustal Omega with amino acid sequence of the putative copper containing protein [40].

### Endodermal spatio-temporal expression pattern

The spatio-temporal endodermal expression pattern of the corresponding genes was checked using the Bio-Analytic Resource database from the AtGenExpress Consortium [41].

### Ionome analysis

The shoot elemental content was measured using Inductively Coupled Plasma Mass Spectrometry (ICP-MS) and the analysis was performed as described [42]. Briefly, shoots of 2-week-old plants grown on agar plates were harvested into Pyrex test tubes (16 x 100 mm) to be then dried at 88 ^o^C for 20h. After weighing the appropriate number of samples (these weights were used to calculate the weights of rest of the sample), the trace metal grade nitric acid Primar Plus (Fisher Chemicals) spiked with indium internal standard was added to the tubes (1 mL per tube). The samples were then digested in a dry block heater (DigiPREP MS, SCP Science; QMX Laboratories, Essex, UK) at 115°C for 4 hours. The digested samples were diluted to 10 mL with 18.2 MΩcm Milli-Q Direct water (Merck Millipore). Elemental analysis was performed using an ICP-MS, PerkinElmer NexION 2000 equipped with Elemental Scientific Inc. autosampler, in the collision mode (He). Twenty-three elements (Li, B, Na, Mg, P, S, K, Ca, Mn, Fe, Co, Ni, Cu, Zn, As, Rb, Sr, Mo, Cd) were monitored. Liquid reference material composed of pooled samples was prepared before the beginning of sample run and was used throughout the whole samples run. It was run after every ninth sample to correct for variation within ICP-MS analysis run. The calibration standards (with indium internal standard and blanks) were prepared from single element standards (Inorganic Ventures;

Essex Scientific Laboratory Supplies Ltd, Essex, UK) solutions. Sample concentrations were calculated using external calibration method within the instrument software. Further data processing was performed using Microsoft Excel spreadsheet. Principal component and heatmap were generated using ClustVis [43].

### Histological staining of roots

Propidium iodide staining was performed as previously described [5]. Root clearing and staining with Calcofluor White, Basic Fuchsin and Auramine-O was performed according to [44]. Fluorol Yellow 088 staining of the suberin was performed as previously described [4,6].

### UCC1 immunolocalization

Custom affinity-purified polyclonal anti-UCC1 antibodies were produced by Genscript, USA and were used as a primary antibody. Anti-UCC1 antibodies were generated in rabbits against a recombinant truncated UCC1 (TDHTIGGPSGWTVGASLRTWAAGQTFAVGDNLVFSYPAAFHD VVEVTKPEFDSCQAVKPLIT-FANGNSLVPLTTPGKRYFICGMPGHCSQGMKLEVNVVPTATVAPTA) produced in *E. coli*. UCC1-immunolocalization was performed according to [45]. 6-day-old seedlings were vacuum infiltrated and fixed with 2% formaldehyde in MTSB buffer (microtubule-stabilizing buffer) supplemented with 0.1 % Triton for 1h. A stock solution of 2x MTSB was prepared with: 15 g PIPES, 1.90 g EGTA, 1.22 g MgSO4o7H2O and 2.5 g KOH and dissolved in a total volume of 500 mL water at pH 7.0 (adjusted with 10 M KOH). Seedlings were washed twice in water that was then replaced by 100% methanol. Methanol content in the wash was gradually decreased until its final concentration reached ~20 %. Seedlings were then washed twice in water and incubated in a cell wall digestion solution (0.2% Driselase, 0.15% Driselase in 2mM MES, pH 5.0) for 30 min at 37°C. After washing with MTSB buffer, the seedlings were incubated in a solution containing 3% IGEPAL CA-630, 10% DMSO in MTSB buffer for 15min at 37°C in order to permeabilise the cell membranes. Seedling were then washed 4 times with MTSB buffer and blocked using a blocking solution containing 2% albumin fraction V BSA in MTSB buffer for 20 min at room temperature. The primary anti-UCC1 antibody was diluted (1/500) in the blocking solution and added to the seedlings for 1h incubation at37°C.Then seedlings were washed twice with MTSB buffer and incubated with the secondary antibody goat anti-Rabbit IgG DyLight 633 (Invitrogen) in MTSB buffer for 1h at 37°C. After washing three times in MTSB buffer, the seedlings were mounted on microscopic slides in MTSB buffer for the microscopy analysis.

### Microscopy

Laser scanning confocal microscopy was performed with a Zeiss LSM500 and a Leica SP8. Structure illumination microscopy was performed with a Zeiss PS1 Super Resolution Microscope. The following excitation and emission detection settings were applied: Calcofluor White, 405 nm/425-475 nm; GFP, 488 nm/500-550 nm; Fluorol yellow 088, 488 nm/500-550 nm; Auramine-O 488nm/505-530nm; propidium iodide, 561 nm/600-620 nm; Basic Fuchsin, 561 nm/570-650 nm; mCherry, 561 nm/ 570-620 nm; Dylight 633, 633 nm/640-670 nm.

### Pixel quantification across the Casparian strip

Images of CS surface view were analysed for *CASP1-GFP*, lignin (Basic fuchsin*/ mCherry-UCC1* and DyLight 633 using Fiji [46]. A segmented line with a width of 3.8μm was traced along the central part of the CS. This selection was then straighten using the “Straighten…” function. A profile of pixels intensity along the *y*-axis was then generated from the pericycle side toward the cortical side. Each replicate represents the average value of pixels intensity across 13 μm approximately and it was generated from one picture. The pixels intensity values were normalised to compare the profile of pixel intensity across the different pictures. The intensity values were scaled from 0 to 1 using the following formula: (*x*-*x*_MIN_)/(*x*_MAX_-*x*_MIN_). In order to obtain a normalisation based solely on CS signal and not on ectopic lignification, the values corresponding to ectopic lignification were intentionally omitted, in the cases the fluorescence signal intensity coming from ectopic lignification (corner of the endodermal cells) was higher than the one coming from the CS itself in the single picture. The profiles of normalised pixel intensity were plotted using PlotTwist [47] and RStudio. Statistical analysis was performed on normalised data in the ranges defined in the figures.

### Timing of expression and localization of CASP1 and UCC1

The timing of CASP1 and UCC1 accumulation and localization at the CS domain were quantified as number of cells after the onset of elongation as previously described [12,31]. The cell number for the endodermal accumulation of CASP1 and UCC1 was determined in the first cell after the onset of elongation where GFP or mCherry signal were detectable using confocal microscopy.

## Acknowledgements

We thank Dr Robert Markus (Super Resolution Microscopy Facility, University of Nottingham), the Microscopy and Histology Facility of the University of Aberdeen and John Danku for ICP-MS analysis. We thank Dr Inês Barbosa and Prof Niko Geldner for critical reading of the manuscript. This work was supported by grants from the UK Biotechnology and Biological Sciences Research Council Grant (grant no. BB/N023927/1 to D.E.S.), the Coordinating Action in Plant Sciences Promoting sustainable collaboration in plant sciences (grant no. ERACAPS13.089_RootBarriers to DES) and a Nottingham Research Fellowship to GC.

